# Distill-Mol: Learning unified molecular representations via cross-modal fusion and hierarchical geometric distillation

**DOI:** 10.64898/2026.11.17.767226

**Authors:** Zihe Liu, Zhonghan Wu, Zhuo Chen, Xin Gao, Bin Yu

**Affiliations:** School of Data Science, Qingdao University of Science and Technology, Qingdao, 266061, China; Computational Bioscience Research Center (CBRC), Computer, Electrical and Mathematical Sciences and Engineering Division, King Abdullah University of Science and Technology (KAUST), Thuwal, 23955, Saudi Arabia; School of Artificial Intelligence and Data Science, University of Science and Technology of China, Hefei, 230027, China

**Keywords:** Molecular representation learning, Multimodal learning, Knowledge distillation, Geometric learning, Molecular property prediction

## Abstract

Molecular representation learning aims to link molecular structures to downstream tasks in drug discovery and materials design. Molecules contain multi-level information, including semantic, topological, and geometric features. However, existing methods often rely on single-modality representations or shallow fusion strategies, limiting their ability to capture complex structure-property relationships. To address this issue, this paper proposes Distill-Mol, a multimodal molecular representation learning framework that formulates molecular representation learning as a language-structure-physics (LSP) collaborative modeling problem. Distill-Mol aligns SMILES semantics, molecular topology, and three-dimensional geometric information in a shared representation space and transfers geometric knowledge from a teacher trained with conformation angle noise to a student trained with coordinate Gaussian noise, enhancing geometric consistency and structural interpretability. In the fusion stage, cross-modal interaction and adaptive gating integrate complementary semantic, topological, and geometric information into task-adaptive molecular representations. Experimental results on MoleculeNet and QM9 demonstrate that Distill-Mol achieves competitive performance across diverse molecular prediction tasks. Additional experiments on DTA and DTI benchmarks further evaluate its applicability to biomedical interaction prediction. Ablation studies, perturbation validation, and interpretability analyses show that Distill-Mol learns task-relevant multimodal representations, improving predictive performance, robustness, and structural interpretability.

## 1. Introduction

Molecular representation learning is a fundamental problem in computational chemistry and molecular modeling, playing a crucial role in practical applications such as drug discovery [1, 2] and materials design [3]. It aims to learn informative molecular representations that can support diverse downstream tasks, including molecular property prediction, drug-target affinity prediction, and drug-target interaction prediction. These tasks require accurate modeling of key molecular characteristics such as solubility [4], toxicity [5], and binding affinity [6], without relying on expensive experiments or complex quantum-chemical calculations. Traditional experimental assays and quantum-chemical methods are often costly and time-consuming, making them difficult to apply to large-scale screening of chemical space [7]. Generalization across molecular scaffolds is also a challenge in molecular property prediction, particularly when labeled data are scarce [8]. These applications require molecular representations that capture complementary structural information and distinguish subtle molecular variations associated with changes in properties.

Deep learning methods learn molecular representations from different structural views to support downstream property prediction. Existing methods typically represent molecules from the perspectives of 1D sequences [9], 2D topology [10], or 3D conformations [11, 12]. Specifically, SMILES sequences encode molecular symbolic semantics, molecular graphs describe atom-bond connectivity and topological structures, and 3D conformations further capture spatial geometry and potential physical constraints. Although these methods have achieved notable progress, individual modalities provide different and incomplete views of molecular structures. Learning how to integrate their complementary information is therefore important for obtaining informative molecular representations.

From this perspective, molecular representation learning can be formulated as a language-structure-physics (LSP) collaborative modeling problem. Under the LSP framework, SMILES sequences correspond to the language level by capturing molecular chemical semantics and contextual dependencies; 2D molecular graphs characterize the structure level by describing atomic connectivity and chemical bond relationships; and 3D conformations represent the physics level by encoding spatial geometry and physical constraints. LSP collaborative modeling aims to achieve alignment, complementarity, and fusion among these information levels within a unified framework, providing a more comprehensive characterization of structure-property relationships. However, two challenges remain. First, effectively integrating heterogeneous molecular representations requires capturing complementary information while preserving modality-specific features, as the contributions of different modalities vary across prediction tasks. Second, learning complementary geometric knowledge from different scales of molecular perturbations remains challenging. Rotations around rotatable bonds produce conformation-level variations, whereas coordinate perturbations introduce local geometric changes. How to connect the information provided by these perturbations within a teacher–student learning framework warrants further investigation. These challenges motivate a framework that combines task-adaptive cross-modal integration with geometric knowledge transfer across perturbation scales.

To address these challenges, we propose Distill-Mol, a multimodal molecular representation learning framework based on dynamic cross-modal fusion and geometry-aware distillation. Within the LSP framework, Distill-Mol extracts semantic, topological, and geometric representations from SMILES sequences, 2D molecular graphs, and 3D conformations, respectively. Cross-modal attention captures complementary information among these representations, while modality self-enhancement and adaptive gating preserve modality-specific features and adjust their contributions according to molecular characteristics and downstream tasks. Meanwhile, geometry-aware distillation connects conformation-level variations induced by conformation angle noise (CAN) with local geometric perturbations introduced by coordinate Gaussian noise (CGN). Specifically, the teacher learns from CAN-perturbed conformations and transfers geometric knowledge to the student trained with CGN-perturbed conformations through hierarchical distillation. Together, these mechanisms integrate complementary molecular information and geometric knowledge into task-adaptive representations for molecular property prediction.

Our contributions can be summarized as follows:

- We propose a geometry-aware hierarchical distillation mechanism that transfers geometric knowledge from a teacher trained with CAN-perturbed conformations to a student trained with CGN-perturbed conformations, connecting conformation-level variations with local geometric perturbations.
- We develop a task-aware dynamic cross-modal fusion strategy within a language-structure-physics (LSP) framework. Cross-modal attention and adaptive gating integrate complementary SMILES, topological, and geometric representations while preserving modality-specific information.
- Extensive experiments on MoleculeNet and QM9 demonstrate competitive performance across diverse molecular prediction tasks. Ablation studies, perturbation analyses, fusion-mechanism analyses, and interpretability analyses provide further evidence for the effectiveness of the proposed framework.

## 2. Related Work

### 2.1. Molecular Representation Learning

In recent years, molecular representation learning has made substantial progress, and the research paradigm has gradually shifted from handcrafted molecular descriptors to data-driven deep learning methods. Existing approaches generally model molecules from three perspectives. The first is SMILES-based semantic modeling, which learns chemical language representations using Transformer architectures, such as ChemBERTa [9]. The second is molecular graph-based structural modeling [13], which employs graph neural networks to characterize interatomic interactions and molecular topology [6, 14]. For example, MPMol [15] uses metapaths to capture functional-group information in heterogeneous molecular graphs and applies hierarchical attention to aggregate node- and relation-level information. The third incorporates three-dimensional geometric information [12], where equivariant neural networks are used to learn molecular spatial conformations and geometric relationships [16]. In addition to directly constructing graph neural network models, molecular graph contrastive learning and data augmentation methods have also been introduced to improve the robustness of molecular representations. For example, LEMON [17] uses line graphs to model bond-level information and applies contrastive objectives to learn molecular graph representations.

Although these methods have achieved notable progress in molecular property prediction, different representation paradigms usually focus on specific levels of molecular information. Sequence-based methods emphasize chemical semantics, graph-based methods focus on topological connectivity, and three-dimensional conformation-based methods mainly capture spatial geometry. Therefore, how to fully exploit these complementary information sources within a unified framework remains an important challenge for complex molecular property prediction [10, 11].

### 2.2. Geometric Self-Supervised Learning and Conformation-Aware Pretraining

Three-dimensional geometric information provides an important complement to molecular topology by describing atomic coordinates, interatomic distances, and bond angles. Equivariant neural networks, such as SE(3)-Transformer [16] and TorchMD-Net [18], incorporate molecular symmetry constraints into geometric representation learning. Sun et al. [19] use tetrahedra as geometric units for self-supervised molecular pretraining. Self-supervised objectives, including contrastive learning [20] and coordinate denoising [21], further enable models to learn from unlabeled molecular conformations. Luo et al. [22] use pseudo force fields and meta-learning to adapt 3D denoising pretraining to downstream property prediction tasks.

Beyond directly learning geometric representations, geometry-informed pretraining also uses 3D information to improve molecular graph representations. GraphMVP [23] exploits the correspondence between 2D topology and 3D geometry through self-supervised pretraining, while 3D Infomax [24] maximizes mutual information between graph and geometric representations. These studies demonstrate the value of incorporating geometric knowledge into molecular graph representation learning.

Building on these studies, we investigate geometric knowledge transfer across perturbation scales. Rotations around rotatable bonds introduce conformation-level variations, whereas coordinate perturbations produce local geometric changes. Distill-Mol connects these perturbations through geometry-aware hierarchical teacher–student distillation, with a teacher trained on CAN-perturbed conformations providing geometric supervision to a student trained on CGN-perturbed conformations.

### 2.3. Multimodal Molecular Fusion and Cross-Modal Knowledge Transfer

Multimodal molecular representation learning has recently attracted increasing attention, aiming to integrate SMILES sequences, 2D topology, 3D conformations, and other molecule-related information to improve molecular representation capability. Existing studies have mainly explored this problem from the perspectives of multi-view alignment, knowledge transfer, and fusion strategies. For example, Nguyen et al. [25] proposed a unified multimodal framework for molecular property prediction, which combines SMILES representations with predicted 3D conformations and uses contrastive learning to enhance consistency across modalities. M2UMol [26] adopts a multimodal-to-unimodal knowledge transfer pretraining strategy, transferring multimodal information into a unified molecular representation to improve downstream generalization. In addition, MolGT [27] integrates multi-source molecular information with a graph Transformer, demonstrating the ability of Transformer architectures to model contextual relationships across modalities. MolMFD [28] further distinguishes modality-shared and modality-specific information through fusion and decoupling mechanisms, thereby improving the discriminative ability and generalization of molecular representations. HMMF-MPP [29] uses task-guided routing in a mixture-of-experts framework to fuse molecular structure and fingerprint representations.

These studies have advanced multimodal molecular representation learning through multi-view alignment, knowledge transfer, and relational fusion. However, most methods focus on integrating molecular features from different sources, while explicit collaboration among language-level semantics, structure-level topology, and physics-level geometry remains underexplored. In contrast, this work reorganizes multimodal molecular representation learning from the perspective of language-structure-physics (LSP) collaborative modeling, and combines geometry-aware distillation with a task-aware dynamic fusion mechanism to improve the consistency, complementarity, and geometric robustness of multimodal representations.

## 3. Method

### 3.1. Overall Framework

The overall framework of Distill-Mol is illustrated in Fig. 1. Distill-Mol is designed for molecular property prediction and formulates molecular representation learning as a language-structure-physics (LSP) collaborative modeling process. Given a molecule, Distill-Mol simultaneously utilizes its 1D SMILES sequence, 2D molecular graph, and 3D conformation to learn molecular representations from three levels: chemical language, topological structure, and spatial geometry.

**Fig. 1.**
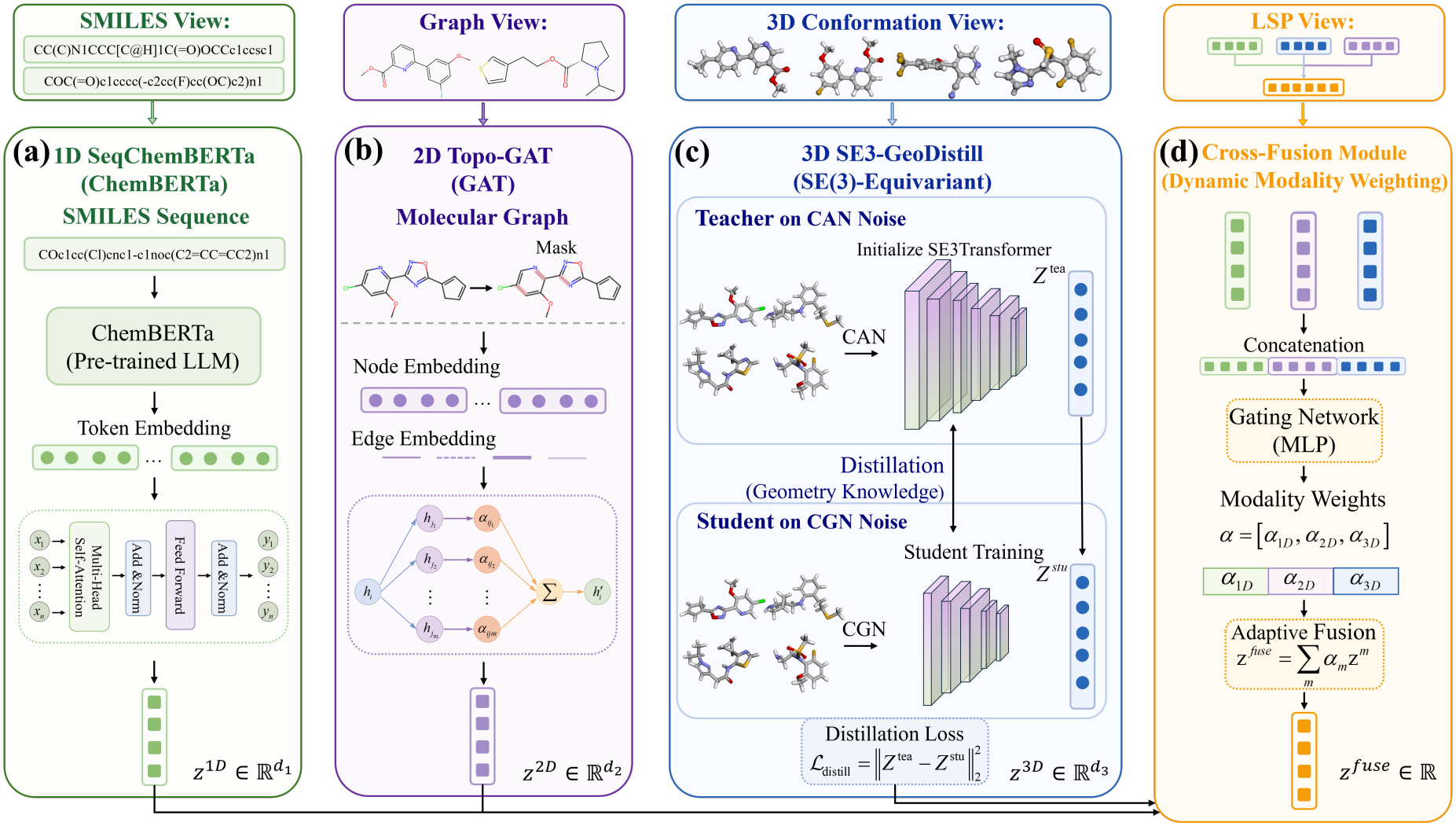
Framework of Distill-Mol. (a) The 1D SeqChemBERTa module encodes SMILES sequences to extract chemical semantic information. (b) The 2D Topo-GAT module models atomic connectivity and local topological structures based on 2D molecular graphs. (c) The 3D SE3-GeoDistill module learns conformation-robust geometric representations through CAN/CGN perturbations and teacher-student distillation. (d) The Cross-Fusion module adaptively integrates 1D, 2D, and 3D representations and feeds the fused representation into the prediction head.

Specifically, the SMILES sequence is fed into the 1D SeqChemBERTa module (Fig. 1a), where the pretrained chemical language model ChemBERTa extracts latent chemical semantic information from the molecular sequence. The two-dimensional molecular graph is processed by the 2D Topo-GAT module (Fig. 1b), in which atoms and chemical bonds are treated as nodes and edges, respectively, while graph attention is used to model atom-level connectivity and local topological patterns. The three-dimensional conformation is further input into the 3D SE3-GeoDistill module (Fig. 1c), which adopts a teacher-student distillation architecture. In this module, conformations perturbed by conformation angle noise (CAN) and coordinate Gaussian noise (CGN) are provided to the teacher and student models, respectively. The former captures conformation-level variations induced by rotatable single bonds, whereas the latter learns stable representations under local geometric perturbations. Through the geometric distillation loss, conformation-level geometric knowledge learned by the teacher model is transferred to the student model, thereby enhancing the geometric consistency and conformational robustness of the 3D representation.

After obtaining the 1D semantic, 2D topological, and 3D geometric representations, these features are fed into a task-aware Cross-Fusion module (Fig. 1d). This module adaptively assigns modality weights according to the dependence of downstream tasks on different modalities, producing a unified molecular representation. Finally, the fused representation is passed to a prediction head for molecular property classification or regression.

### 3.2. Language-Structure-Physics Collaborative Encoder

Within the overall framework, Distill-Mol extracts language, topological, and geometric representations from SMILES sequences, 2D molecular graphs, and 3D conformations, respectively. The three encoders correspond to different information levels in the language-structure-physics (LSP) framework and are used to construct unified multimodal molecular representations, providing the basis for subsequent cross-modal collaborative modeling.

#### 3.2.1. One-Dimensional Language Modeling

The SeqChemBERTa module is designed to perform semantic encoding of molecular SMILES sequences. A SMILES sequence represents molecular composition, connectivity, and local chemical semantics in a linear symbolic form [9]. In this work, SMILES is regarded as a molecular language governed by syntactic rules and enriched with chemical semantics. Accordingly, the pretrained chemical language model ChemBERTa is employed to extract molecular semantic representations. Given the SMILES sequence *S* = (*t*_1_, *t*_2_, …, *t*_*L*_) of a molecule *M*, where *t*_*i*_ denotes the *i*-th token and *L* denotes the sequence length, the sequence is fed into the pretrained language model for contextual encoding, yielding token-level hidden representations:

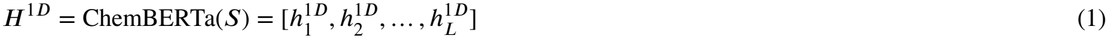

where 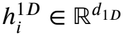 denotes the contextual semantic representation of the *i*-th token, and *d*_1*D*_ is the feature dimension of the 1D semantic space. To obtain a molecule-level semantic representation, the token-level hidden representations are aggregated by a pooling operation:

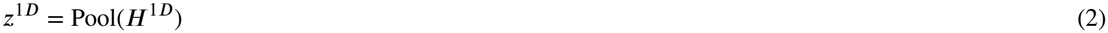

where Pool(*⋅*) denotes the aggregation operation over token-level hidden representations, and *z*^1*D*^ denotes the global semantic representation of the molecule. This representation captures contextual dependencies in the SMILES sequence, thereby encoding high-level information such as functional group combinations, local chemical environments, and potential reaction semantics, and provides semantic priors for subsequent multimodal fusion.

#### 3.2.2. Two-Dimensional Topological Modeling

The Topo-GAT module is designed to encode the topological structure of 2D molecular graphs. A molecular graph explicitly describes atomic connectivity and local topological structures [13, 30]. Given a molecular graph *G* = (*V*, *E*), where *V* denotes the set of atom nodes and *E* denotes the set of chemical bond edges, the initial representation of each atom node *v*_*i*_ *∈ V* is obtained by encoding atomic attributes such as atom type, valence electron number, aromaticity, and hybridization state. For each chemical bond *e*_*ij*_ *∈ E*, its edge representation is obtained from bond-related attributes such as bond type, conjugation status, and ring membership:

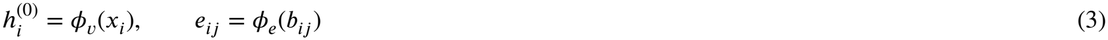

where *x*_*i*_ denotes atom attributes, *b*_*ij*_ denotes chemical bond attributes, and *ϕ*_*v*_(*⋅*) and *ϕ*_*e*_(*⋅*) are learnable mapping functions. To model the 2D topological structure of a molecule, we adopt the graph attention network Topo-GAT to encode the graph structure. At the *l*-th layer, the attention weight from node *i* to its neighboring node *j* is defined as:

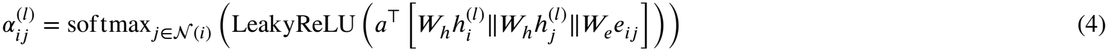

where *N* (*i*) denotes the neighbor set of node *i, ‖*denotes vector concatenation, and 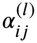 denotes the attention weight from node *i* to neighboring node *j*. The node representation is updated as:

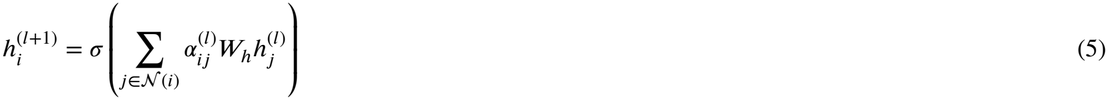

After *L*_*g*_ layers of graph attention encoding, node-level topological representations are obtained. A graph-level readout function is then used to aggregate them into a molecule-level 2D topological representation:

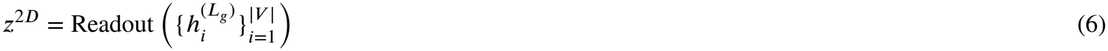

where 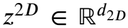 denotes the 2D topological representation of the molecule, which encodes atomic connectivity, local structural patterns, and key topological substructure features.

#### 3.2.3. Three-Dimensional Geometric Modeling

The SE3-GeoDistill module is designed to learn geometric representations from 3D molecular conformations. Compared with 2D molecular topology, 3D conformations provide explicit spatial geometric information, including interatomic distances, conformational shapes, and local geometric relationships that are difficult to directly capture from 2D topology alone [18, 31]. Given a molecular graph *G* and its atomic coordinates *X ∈* ℝ^*N*×3^, we employ an SE(3)-equivariant geometric encoder to model the 3D molecular structure:

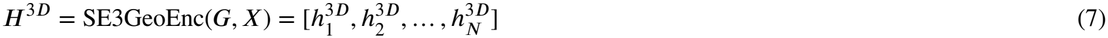

where 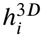 denotes the geometric representation of the *i*-th atom. SE(3)-equivariant networks preserve the consistency of geometric representations under 3D rotations and translations, making them suitable for modeling spatial relationships and local geometric structures in molecular conformations. Furthermore, a molecule-level 3D geometric representation is obtained through an atom-level pooling function:

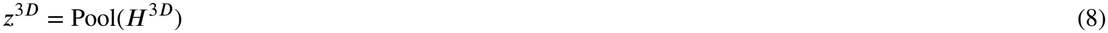

where 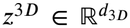 denotes the 3D geometric representation of the molecule, which is used to characterize spatial conformations, interatomic geometric relationships, and local structural patterns. Based on this geometric encoder, the subsequent geometry-aware distillation further enhances the conformational robustness and geometric consistency of 3D representations through a teacher-student mechanism under CAN/CGN perturbations.

### 3.3. Geometry-Aware Distillation

#### 3.3.1. Conformation Perturbation Design

Three-dimensional conformations provide spatial geometric information of molecules, but directly modeling a single conformation is often insufficient to fully characterize molecular variations in realistic conformational space [31, 32]. In particular, for molecules containing rotatable single bonds, conformational changes involve both local perturbations around equilibrium conformations and larger-scale conformational differences induced by torsional variations [33]. Subsequent energy distribution analysis shows that CGN-generated conformations are mainly concentrated in low-energy regions, whereas CAN-generated conformations span a broader range of conformational variations. These results suggest that the two perturbation strategies are complementary in terms of perturbation scale and energy distribution. Accordingly, the teacher model provides larger-scale conformational geometric knowledge, while the student model learns robust geometric representations under locally stable perturbations, thereby establishing a reasonable foundation for subsequent geometric distillation.

To address this issue, we design a geometry-aware teacher-student distillation mechanism, as illustrated in Fig. 2. Specifically, conformation angle noise (CAN) perturbs torsion angles around rotatable single bonds to generate conformation-level variations and is used as the input to the teacher model, while coordinate Gaussian noise (CGN) introduces local continuous perturbations to atomic coordinates and is used as the input to the student model. In this way, CAN and CGN provide complementary geometric supervision at different perturbation scales, enabling cross-scale geometric knowledge transfer for 3D molecular representation learning [34, 35].

**Fig. 2.**
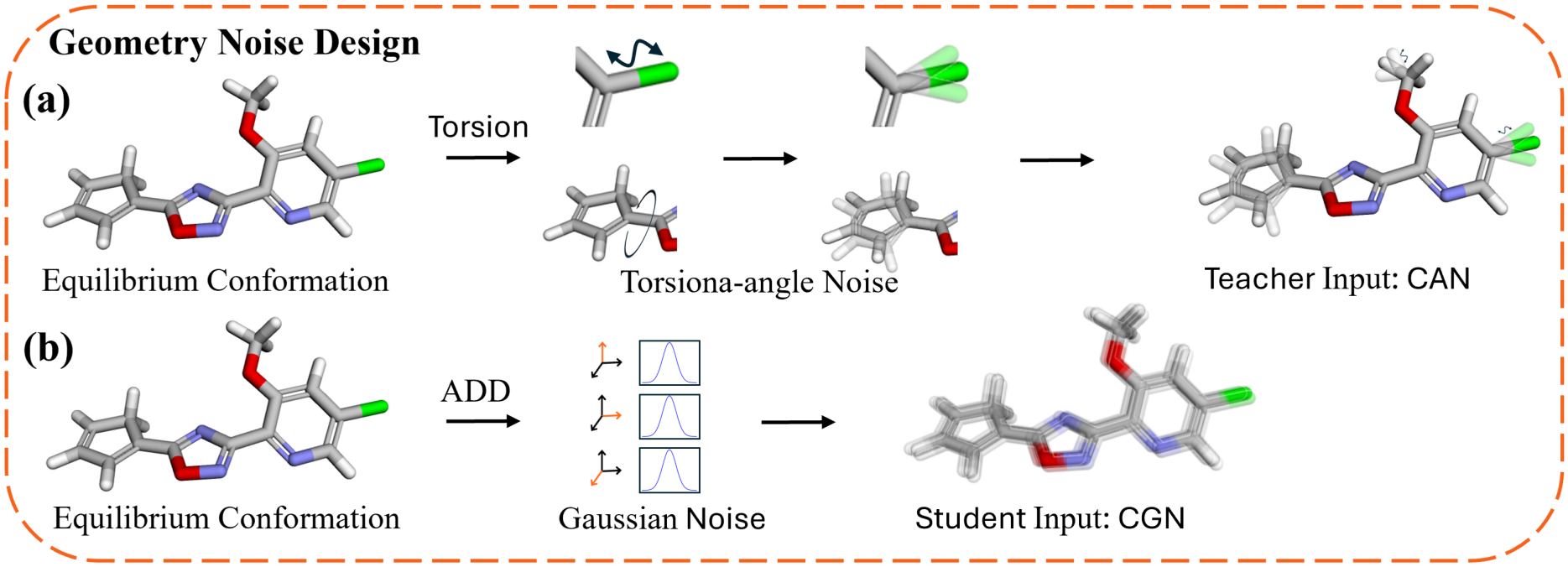
Geometry-aware noise design for teacher-student geometric distillation. CAN generates conformation-level variations through torsion-angle perturbations and is fed into the teacher model, whereas CGN simulates local geometric perturbations via coordinate Gaussian noise and is fed into the student model.

Given the original three-dimensional conformation *X ∈* ℝ^*N*×3^ of a molecule, we first construct conformation angle noise CAN. Let ℜ denote the set of rotatable single bonds in the molecule. For each rotatable single bond *r ∈* ℜ, a torsional perturbation is sampled as:

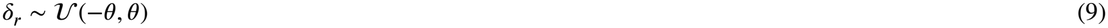

where *θ* denotes the maximum perturbation angle. Based on the sampled torsional perturbation set *δ* = {*δ*_*r*_}_*r∈*ℜ_, the CAN-perturbed conformation is generated through the conformation transformation operator *T*_*δ*_(*⋅*):

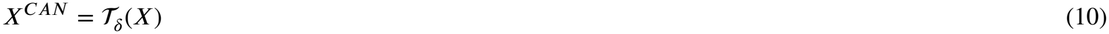

This process simulates structural variations of the molecule in conformational space by changing the local torsional states around rotatable single bonds, thereby providing conformation-level geometric perturbation information for the model.

Unlike CAN, CGN directly acts on atomic coordinates and is used to simulate local continuous perturbations around the original conformation:

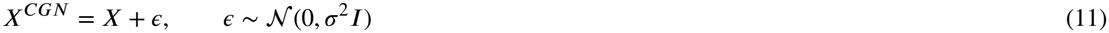

where ε *∈* ℝ ^*N*×3^ denotes the 3D coordinate noise matrix applied to all atoms, and σ controls the intensity of the Gaussian perturbation. CAN focuses more on conformation-level differences induced by torsional variations, whereas CGN mainly characterizes local geometric perturbations around the original conformation. These two perturbations complement each other, enabling the model to perceive both larger-scale conformational changes and local spatial perturbations.

#### 3.3.2. Teacher-Student Geometric Distillation

Based on the perturbation construction described above, we build a teacher-student geometric distillation framework. The teacher model receives CAN-perturbed conformations to capture conformation-level geometric variations, whereas the student model receives CGN-perturbed conformations to learn stable geometric representations under local perturbations. Both models adopt an SE(3)-equivariant geometric encoder:

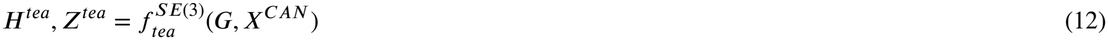

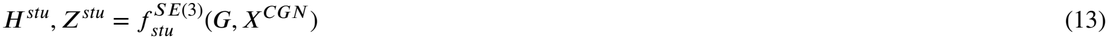

where *H*^*tea*^ and *H*^*stu*^ denote the atom-level geometric representations produced by the teacher and student models, respectively, while *Z*^*tea*^ and *Z*^*stu*^ denote the corresponding molecule-level geometric representations. By transferring conformation-level geometric knowledge learned by the teacher under CAN perturbations to the student model, the proposed framework enables the student to acquire more robust 3D molecular representations from locally perturbed conformations.

#### 3.3.3. Distillation Objective

To achieve cross-scale geometric knowledge transfer, we impose consistency constraints between the teacher and student models on atom-level hidden representations. Specifically, the student and teacher models output multi-layer atom-level feature representations, and the atom features from the *l*-th student layer are aligned with those from the corresponding teacher layer. When the teacher and student models have different numbers of layers, a uniform layer mapping strategy is used to select the corresponding teacher layer. When their hidden dimensions are inconsistent, a feature alignment layer is used to map student features into the teacher feature space.

The atom-level inter-layer distillation loss is defined as:

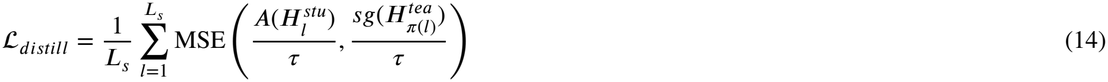

where 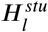 denotes the atom-level features output by the *l*-th layer of the student model, 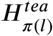 denotes the atom-level features of the corresponding teacher layer, *A*(*⋅*) denotes the feature alignment mapping, *π*(*l*) denotes the mapping relationship from the student layer to the teacher layer, *τ* is the temperature coefficient, and *sg*(*⋅*) denotes the stop-gradient operation. This loss encourages the student model, under CGN local perturbation inputs, to progressively fit the atom-level geometric representations learned by the teacher model under CAN conformational perturbations.

##### Algorithm 1

Dynamic Cross-Modal Fusion

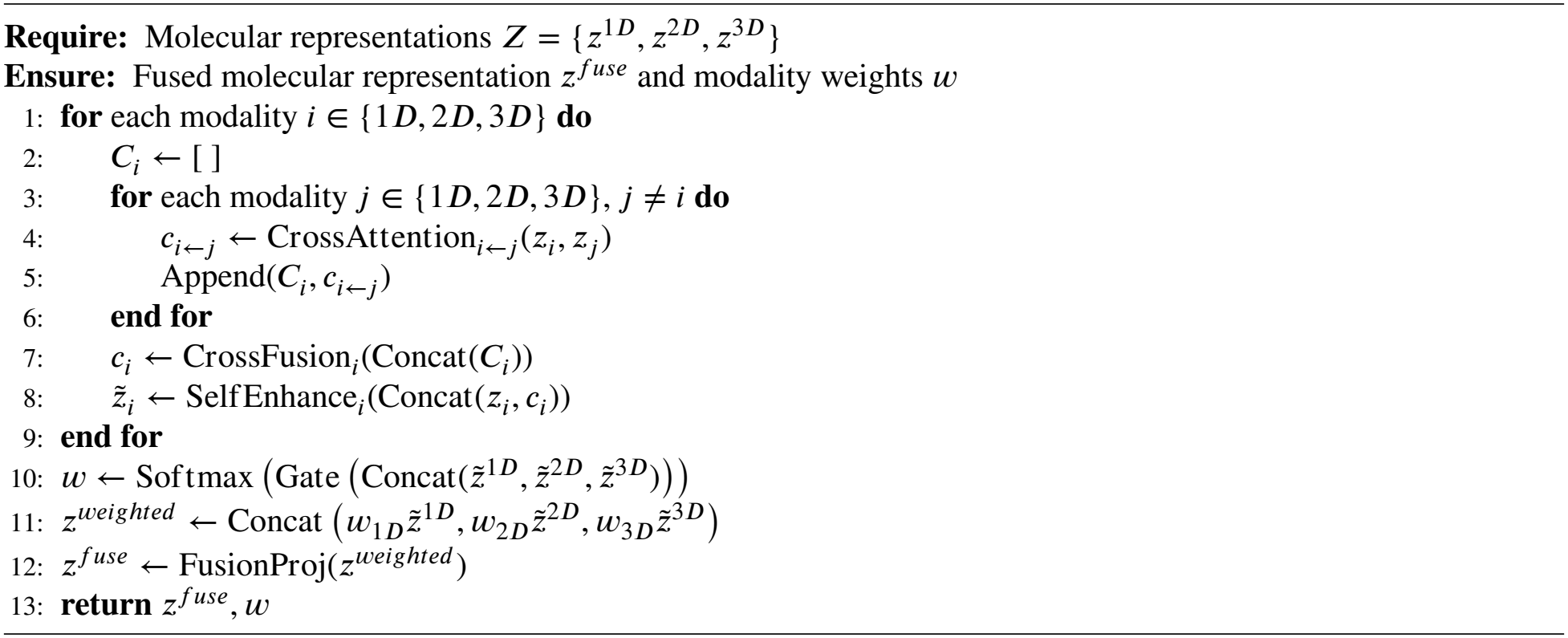

In addition to the distillation loss, the student model employs a coordinate denoising loss ℒ_*denoise*_ to enhance local geometric recovery and an auxiliary centroid prediction loss ℒ_*aux*_ to stabilize graph-level representation learning. Therefore, the overall objective in the pretraining stage is defined as:

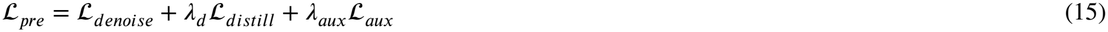

where *λ*_*d*_ is the weight of the distillation loss, and *λ*_*aux*_ is the weight of the auxiliary loss. Through the above objective, the student model not only learns local coordinate denoising capability, but also absorbs atom-level geometric knowledge acquired by the teacher model under conformational perturbations, thereby enhancing the robustness and geometric consistency of 3D representations against conformational variations.

### 3.4. Cross-Modal Fusion and Prediction

The Cross-Fusion module is designed to dynamically integrate the 1D language, 2D topological, and 3D geometric representations of a molecule. After the modality-specific encoding process, each molecule is represented by *z*^1*D*^, *z*^2*D*^, and *z*^3*D*^. Since different downstream tasks may depend on different molecular information, directly concatenating these representations may introduce redundant features and cannot sufficiently model cross-modal interactions. To address this issue, we design a task-aware dynamic cross-modal fusion strategy that combines cross-modal attention, self-enhancement, and adaptive gating to capture complementary structure-semantics-geometry information.

The overall fusion process is shown in Algorithm 1. First, for each modality *i ∈* {1*D*, 2*D*, 3*D*}, cross-modal attention is used to aggregate information from the other modalities. Specifically, the representation of modality *i* attends to the representation of modality *j*, where *j* ≠ *i*, producing the cross-modal context *c*_*i*←*j*_. The contexts from different source modalities are then concatenated and passed through a cross-fusion layer to obtain the integrated context representation *c*_*i*_. Subsequently, *c*_*i*_ is combined with the original modality representation *z*_*i*_ through a self-enhancement module, yielding the enhanced modality representation 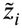.

After obtaining the enhanced representations 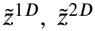, and 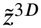, a gating network is employed to adaptively estimate the contribution of each modality:

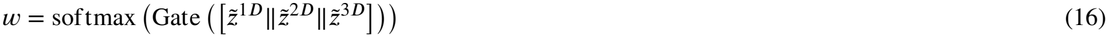

where *w* = [*w*_1*D*_, *w*_2*D*_, *w*_3*D*_] denotes the modality weights, ∑_*m*_ *w*_*m*_ = 1, and ‖ denotes vector concatenation. The gating mechanism enables the model to dynamically adjust the importance of different modalities according to the input molecule and downstream task.

Based on the learned modality weights, the weighted modality representations are concatenated and projected into a unified molecular representation:

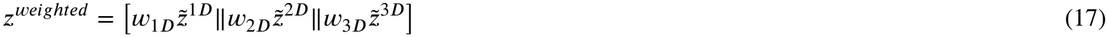

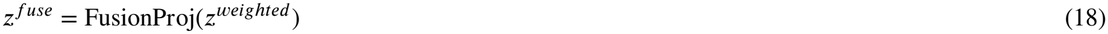

Compared with simple concatenation or fixed weighting, this design preserves modality-specific information while enhancing cross-modal complementarity, thereby reducing information redundancy and improving the collaborative representation of heterogeneous molecular features.

Finally, the fused representation is fed into a task prediction head:

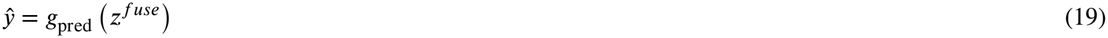

where *g*_pred_(*⋅*) denotes a multilayer perceptron. For classification tasks, the model is optimized using the cross-entropy loss, whereas the mean squared error loss is adopted for regression tasks. Through this dynamic fusion mechanism, Distill-Mol can generate a unified molecular representation that adaptively integrates chemical semantics, molecular topology, and spatial geometry for downstream molecular prediction.

## 4. Experiments

This section systematically evaluates the effectiveness of Distill-Mol on molecular representation learning tasks. We first introduce the experimental datasets, training configurations, and baseline methods. Then, we compare the performance of Distill-Mol with representative baselines on MoleculeNet and QM9. Finally, we validate the proposed language-structure-physics (LSP) collaborative learning framework through analyses of modality complementarity, geometric perturbation, interpretability, and biomedical interaction generalization.

### 4.1. Experimental Setup

#### 4.1.1. Datasets

To comprehensively evaluate Distill-Mol, experiments were conducted on both pretraining datasets and down-stream molecular property prediction benchmark datasets. In the pretraining stage, unlabeled small molecules from the ZINC dataset [36] were used as pretraining data to learn general molecular representations. The ZINC dataset contains a large number of structurally diverse small molecules, providing rich samples for learning molecular semantics, topological structures, and 3D geometric information.

For downstream evaluation, we mainly used two widely adopted molecular learning benchmark datasets, MoleculeNet [37] and QM9 [38]. MoleculeNet covers various tasks, including bioactivity, toxicity, and physicochemical property prediction. We selected BACE, BBBP, ClinTox, Tox21, ToxCast, MUV, SIDER, and HIV as classification tasks, and ESOL, FreeSolv, and Lipophilicity as regression tasks. QM9 contains various quantum-chemical properties and was used to evaluate the model’s ability to capture 3D geometric structures and physical information. In addition, Activity Cliff experiments were constructed based on ChEMBL [39] to further verify the model’s sensitivity to finegrained structural changes.

For MoleculeNet classification tasks, ROC-AUC was used as the evaluation metric. RMSE was adopted for MoleculeNet and Activity Cliff regression tasks, while MAE was used for QM9 quantum property prediction tasks. Data splits were performed using the scaffold split strategy to evaluate model performance under scaffold out-of-distribution scenarios.

#### 4.1.2. Experimental Configuration

All experiments were conducted on an NVIDIA RTX 3090 GPU with 24 GB memory. Distill-Mol was trained using the AdamW optimizer, a learning rate of 2 × 10^−5^, a weight decay of 0.01, a batch size of 32, and 30 training epochs. All experiments were independently repeated using 10 predefined random seeds. The mean and standard deviation across the 10 runs were reported as the final results.

The dataset split was performed using the Scaffold strategy. The Scaffold split was used to construct out-of-distribution test scenarios based on molecular scaffolds, and a Balanced Scaffold strategy was adopted to more rigorously evaluate the generalization ability of the model. During training, the fusion layer dimension was set to 512, and the dropout rate was set to 0.2 to mitigate overfitting and improve model generalization.

#### 4.1.3. Baseline

We compare Distill-Mol with representative baselines using the evaluation protocols adopted for each benchmark. Baseline results were taken from the corresponding publications when available; otherwise, the methods were reproduced using their official implementations. For the reproduced baselines, the same dataset splits and evaluation metrics were used, while model-specific hyperparameters followed the recommended settings in the original studies.

- **ChemBERTa-2** [9]: A Transformer-based pretrained model for SMILES sequences, which treats SMILES as a chemical language and learns contextual semantic representations from large-scale unlabeled molecular sequences.
- **Transformer-M** [40]: Graph Transformer-based molecular representation learning method encoding both 2D topology and 3D structure to capture long-range dependencies and multi-structure information.
- **TorchMD-Net** [18]: SE(3)-equivariant Transformer for 3D molecular modeling, directly leveraging atom types and 3D coordinates to model spatial relationships, suitable for geometry-sensitive property prediction.
- **SE(3)-Transformer** [16]: Attention network equivariant to 3D rotation and translation, effectively modeling spatial geometry in molecular conformations; a representative 3D equivariant model.
- **Chemprop** [41]: Directed message-passing neural network (D-MPNN) for molecular property prediction, propagating information along chemical bonds to capture local chemical environments and atomic interactions.
- **Attentive FP** [42]: Graph attention-based molecular representation method that adaptively focuses on key atoms and substructures relevant to target properties during message passing and readout.
- **MolCLR** [43]: Graph contrastive learning approach using atom masking, bond deletion, and subgraph removal to generate positive views and learn generalizable graph representations.
- **InfoGraph** [44]: Maximizes mutual information between graph-level representations and local node, edge, and subgraph representations to learn molecular embeddings capturing multi-scale structural information.
- **JOAO** [45]: Automated graph contrastive learning method introducing joint augmentation optimization in the GraphCL framework, adaptively selecting optimal augmentations to reduce uncertainty from manual choices.
- **3D Infomax** [24]: 3D molecular representation learning via self-supervised contrastive learning, maximizing mutual information between 2D graph and 3D geometric representations.
- **Mole-BERT** [46]: A BERT-inspired method that learns substructure and semantic representations from unlabeled molecules for downstream property prediction.
- **GraphMVP** [23]: 2D-3D multi-view molecular pretraining method, modeling consistency between 2D topology and 3D geometry to enhance 2D graph encoder representations.
- **MolGT** [27]: Graph Transformer-based multi-modal pretraining method, integrating 2D topology and 3D geometric information, with node-level and graph-level pretraining tasks.

### 4.2. Results

This section reports the performance of Distill-Mol on MoleculeNet and QM9 benchmark datasets and analyzes the effectiveness of the proposed multimodal collaborative modeling strategy. Supplementary DTA and DTI results are summarized in Table S.2 of the Supplementary Material to further evaluate the transferability of the learned molecular representations.

#### 4.2.1. Performance on MoleculeNet

To comprehensively evaluate the performance of Distill-Mol on small-molecule property prediction tasks, we first conduct experiments on the MoleculeNet benchmark dataset. MoleculeNet covers both classification and regression tasks, enabling the evaluation of molecular representation capability from multiple perspectives, including bioactivity, toxicity, safety, and physicochemical properties. For classification tasks, we select BACE, BBBP, ClinTox, Tox21, ToxCast, MUV, SIDER, and HIV, and use ROC-AUC as the evaluation metric, where a higher value indicates better classification performance. For regression tasks, we select ESOL, FreeSolv, and Lipophilicity, and use RMSE as the evaluation metric, where a lower value indicates smaller prediction error. The performance comparisons between Distill-Mol and representative baseline methods on MoleculeNet classification and regression tasks are shown in Table 1 and Table 2 respectively.

**Table 1.**
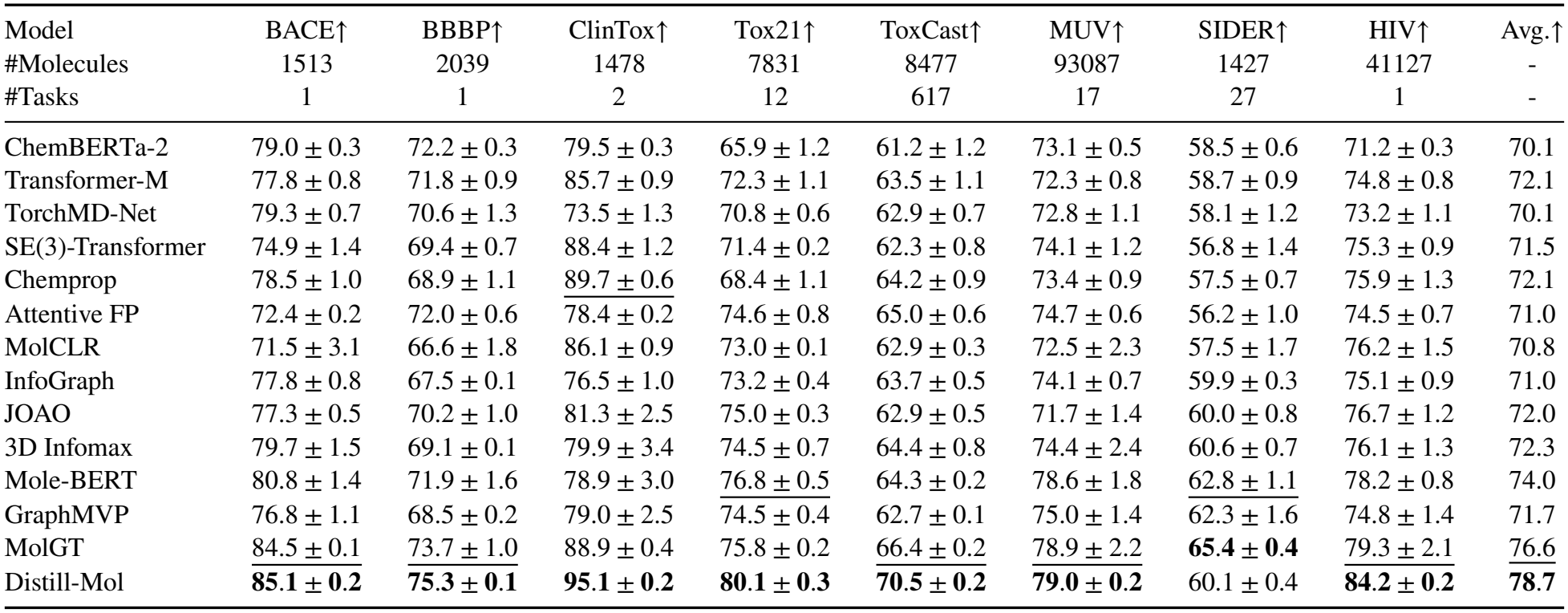
Performance comparison on MoleculeNet small-molecule property prediction tasks. The reported results are mean ROC-AUC ± standard deviation, where higher values indicate better performance. The best results are highlighted in bold, and the second-best results are underlined.

| Model | BACE↑ | BBBP↑ | ClinTox↑ | Tox21↑ | ToxCast↑ | MUV↑ | SIDER↑ | HIV↑ | Avg.↑ |
|---|---|---|---|---|---|---|---|---|---|
| #Molecules | 1513 | 2039 | 1478 | 7831 | 8477 | 93087 | 1427 | 41127 | - |
| #Tasks | 1 | 1 | 2 | 12 | 617 | 17 | 27 | 1 | - |
| ChemBERTa-2 | 79.0 ± 0.3 | 72.2 ± 0.3 | 79.5 ± 0.3 | 65.9 ± 1.2 | 61.2 ± 1.2 | 73.1 ± 0.5 | 58.5 ± 0.6 | 71.2 ± 0.3 | 70.1 |
| Transformer-M | 77.8 ± 0.8 | 71.8 ± 0.9 | 85.7 ± 0.9 | 72.3 ± 1.1 | 63.5 ± 1.1 | 72.3 ± 0.8 | 58.7 ± 0.9 | 74.8 ± 0.8 | 72.1 |
| TorchMD-Net | 79.3 ± 0.7 | 70.6 ± 1.3 | 73.5 ± 1.3 | 70.8 ± 0.6 | 62.9 ± 0.7 | 72.8 ± 1.1 | 58.1 ± 1.2 | 73.2 ± 1.1 | 70.1 |
| SE(3)-Transformer | 74.9 ± 1.4 | 69.4 ± 0.7 | 88.4 ± 1.2 | 71.4 ± 0.2 | 62.3 ± 0.8 | 74.1 ± 1.2 | 56.8 ± 1.4 | 75.3 ± 0.9 | 71.5 |
| Chemprop | 78.5 ± 1.0 | 68.9 ± 1.1 | 89.7 ± 0.6 | 68.4 ± 1.1 | 64.2 ± 0.9 | 73.4 ± 0.9 | 57.5 ± 0.7 | 75.9 ± 1.3 | 72.1 |
| Attentive FP | 72.4 ± 0.2 | 72.0 ± 0.6 | 78.4 ± 0.2 | 74.6 ± 0.8 | 65.0 ± 0.6 | 74.7 ± 0.6 | 56.2 ± 1.0 | 74.5 ± 0.7 | 71.0 |
| MolCLR | 71.5 ± 3.1 | 66.6 ± 1.8 | 86.1 ± 0.9 | 73.0 ± 0.1 | 62.9 ± 0.3 | 72.5 ± 2.3 | 57.5 ± 1.7 | 76.2 ± 1.5 | 70.8 |
| InfoGraph | 77.8 ± 0.8 | 67.5 ± 0.1 | 76.5 ± 1.0 | 73.2 ± 0.4 | 63.7 ± 0.5 | 74.1 ± 0.7 | 59.9 ± 0.3 | 75.1 ± 0.9 | 71.0 |
| JOAO | 77.3 ± 0.5 | 70.2 ± 1.0 | 81.3 ± 2.5 | 75.0 ± 0.3 | 62.9 ± 0.5 | 71.7 ± 1.4 | 60.0 ± 0.8 | 76.7 ± 1.2 | 72.0 |
| 3D Infomax | 79.7 ± 1.5 | 69.1 ± 0.1 | 79.9 ± 3.4 | 74.5 ± 0.7 | 64.4 ± 0.8 | 74.4 ± 2.4 | 60.6 ± 0.7 | 76.1 ± 1.3 | 72.3 |
| Mole-BERT | 80.8 ± 1.4 | 71.9 ± 1.6 | 78.9 ± 3.0 | 76.8 ± 0.5 | 64.3 ± 0.2 | 78.6 ± 1.8 | 62.8 ± 1.1 | 78.2 ± 0.8 | 74.0 |
| GraphMVP | 76.8 ± 1.1 | 68.5 ± 0.2 | 79.0 ± 2.5 | 74.5 ± 0.4 | 62.7 ± 0.1 | 75.0 ± 1.4 | 62.3 ± 1.6 | 74.8 ± 1.4 | 71.7 |
| MolGT | 84.5 ± 0.1 | 73.7 ± 1.0 | 88.9 ± 0.4 | 75.8 ± 0.2 | 66.4 ± 0.2 | 78.9 ± 2.2 | **65.4 ± 0.4** | 79.3 ± 2.1 | 76.6 |
| Distill-Mol | **85.1 ± 0.2** | **75.3 ± 0.1** | **95.1 ± 0.2** | **80.1 ± 0.3** | **70.5 ± 0.2** | **79.0 ± 0.2** | 60.1 ± 0.4 | **84.2 ± 0.2** | **78.7** |

**Table 2.** Performance comparison on MoleculeNet small-molecule property regression tasks. The reported results are mean RMSE ± standard deviation, where lower values indicate better performance. The best results are highlighted in bold, and the second-best results are underlined.

| Model | ESOL | FreeSolv | Lipophilicity |
| #Molecules | 1128 | 642 | 4200 |
| #Tasks | 1 | 1 | 1 |
|---|---|---|---|
| ChemBERTa-2 | 0.912 ± 0.041 | 2.185 ± 0.196 | 0.742 ± 0.018 |
| Transformer-M | 0.864 ± 0.035 | 2.032 ± 0.172 | 0.718 ± 0.021 |
| TorchMD-Net | 0.881 ± 0.038 | 2.114 ± 0.185 | 0.731 ± 0.019 |
| SE(3)-Transformer | 0.856 ± 0.032 | 2.067 ± 0.161 | 0.726 ± 0.020 |
| Chemprop | 0.936 ± 0.044 | 2.248 ± 0.203 | 0.705 ± 0.017 |
| Attentive FP | 0.948 ± 0.047 | 2.296 ± 0.214 | 0.698 ± 0.016 |
| 3D Infomax | 0.894 ± 0.028 | 2.337 ± 0.227 | 0.695 ± 0.012 |
| Mole-BERT | 1.015 ± 0.030 | - | **0.676 ± 0.017** |
| GraphMVP | 1.029 ± 0.033 | - | 0.682 ± 0.010 |
| MolGT | 0.839 ± 0.006 | - | 0.788 ± 0.013 |
| Distill-Mol | **0.793 ± 0.002** | **1.735 ± 0.092** | 0.735 ± 0.024 |

As shown in Table 1 Distill-Mol achieves strong overall performance on MoleculeNet classification tasks, obtaining the best results on 7 out of 8 datasets, including BACE, BBBP, ClinTox, Tox21, ToxCast, MUV, and HIV. In terms of average ROC-AUC, Distill-Mol reaches 78.7, outperforming representative baselines such as MolGT, Mole-BERT, and 3D Infomax, indicating that multimodal collaborative modeling can provide stable improvements across most classification tasks.

This advantage mainly comes from the joint utilization of multi-source molecular information. Compared with sequence-based, graph-based, and 3D geometry-based methods that focus on specific information levels, Distill-Mol integrates 1D semantic, 2D topological, and 3D geometric representations through a dynamic fusion mechanism, enabling adaptive collaboration among different modalities. However, Distill-Mol does not achieve the best result on SIDER, with a ROC-AUC of 60.1, suggesting that multi-label side-effect prediction may require additional side-effect-specific structural priors or label-correlation modeling.

From the regression results in Table 2 Distill-Mol achieves competitive performance on ESOL and FreeSolv, with RMSE values of 0.793 and 1.735, respectively, suggesting its ability to capture molecular features related to water solubility and hydration free energy. In contrast, its performance on Lipophilicity is not optimal, with an RMSE of 0.735, indicating that lipophilicity may depend more on local hydrophobic fragments, molecular fragment distribution, or specific physicochemical descriptors. These results further show that different molecular properties rely on semantic, topological, and geometric information to different extents, highlighting the necessity of task-aware multimodal fusion.

#### 4.2.2. Performance on QM9

To further evaluate the modeling capability of Distill-Mol for geometry-sensitive properties, we conduct quantum-chemical property prediction experiments on the QM9 dataset. QM9 contains multiple quantum-chemical properties related to electronic structure, spatial conformation, and thermodynamic characteristics, enabling the evaluation of the model’s ability to capture 3D geometric information and physical consistency. We select nine tasks, including *µ, α*, HOMO, LUMO, Gap, ZPVE, *U*_0_, *U*, and *H*, and use MAE as the evaluation metric. The performance comparison between Distill-Mol and representative baselines on QM9 is shown in Table 3.

**Table 3.** Performance comparison on QM9 quantum-chemical property prediction tasks. The reported results are mean MAE ± standard deviation, where lower values indicate better performance. The best results are highlighted in bold, and the second-best results are underlined.

| Model | *μ* (D) | *α* (*a*<sub>0</sub><sup>3</sup>) | HOMO | LUMO | Gap | ZPVE | *U*<sub>0</sub> | *U* | *H* |
|---|---|---|---|---|---|---|---|---|---|
| ChemBERTa-2 | 0.015 ± 0.002 | 0.047 ± 0.010 | 16.3 ± 0.24 | 14.2 ± 0.31 | 25.8 ± 0.31 | 1.34 ± 0.12 | 5.61 ± 0.05 | 5.71 ± 0.18 | **5.63 ± 0.06** |
| Chemprop | 0.014 ± 0.003 | 0.042 ± 0.009 | 17.2 ± 0.19 | 14.5 ± 0.29 | 26.1 ± 0.25 | 1.35 ± 0.14 | 5.65 ± 0.08 | 5.65 ± 0.14 | 6.12 ± 0.09 |
| Attentive FP | 0.019 ± 0.002 | 0.042 ± 0.009 | **13.9 ± 0.23** | 14.6 ± 0.33 | 27.7 ± 0.35 | 1.42 ± 0.13 | 5.69 ± 0.09 | 5.72 ± 0.12 | 5.64 ± 0.11 |
| TorchMD-Net | 0.011 ± 0.003 | 0.059 ± 0.011 | 20.3 ± 0.35 | 18.6 ± 0.31 | 36.1 ± 0.29 | 1.84 ± 0.19 | 6.15 ± 0.06 | 6.38 ± 0.09 | 6.16 ± 0.06 |
| SE(3)-Transformer | 0.015 ± 0.002 | 0.046 ± 0.008 | 23.5 ± 0.34 | 19.5 ± 0.25 | 40.2 ± 0.30 | 1.31 ± 0.16 | 6.92 ± 0.07 | 6.99 ± 0.11 | 7.09 ± 0.12 |
| Transformer-M | 0.037 ± 0.004 | 0.041 ± 0.006 | 17.5 ± 0.21 | 16.2 ± 0.24 | 27.4 ± 0.18 | **1.18 ± 0.11** | 9.37 ± 0.08 | 9.41 ± 0.12 | 9.39 ± 0.15 |
| Distill-Mol | **0.010 ± 0.002** | **0.039 ± 0.004** | 15.1 ± 0.21 | **13.6 ± 0.22** | **25.0 ± 0.22** | 1.20 ± 0.09 | **5.51 ± 0.05** | **5.59 ± 0.04** | 5.69 ± 0.04 |

As shown in Table 3 Distill-Mol achieves the lowest MAE on six out of nine QM9 tasks, including *µ, α*, LUMO, Gap, *U*_0_, and *U*. These results demonstrate its strong capability in modeling geometry- and physics-related molecular properties. Compared with models that mainly rely on SMILES sequences or 2D molecular topology, Distill-Mol incorporates 3D geometric information and geometry-aware distillation, enabling it to better capture conformational variations, local spatial relationships, and structural features relevant to quantum-chemical properties. As a result, it achieves more stable performance across most geometry-sensitive tasks. Although Distill-Mol does not achieve the best results on HOMO, ZPVE, and *H*, it still shows competitive overall performance across QM9 tasks. The remaining gaps suggest that some high-precision quantum-chemical properties may depend more heavily on fine-grained electronic structures, vibrational characteristics, or physical constraints. Therefore, incorporating additional physical priors, such as local charges, electron density, molecular orbital information, or molecular force-field constraints, may further improve the prediction of these properties in future work.

### 4.3. LSP Complementarity and Fusion Analysis

To investigate the contributions of language, structure, and physics information in Distill-Mol, we analyze prediction performance, fusion behavior, and representation space. First, ablation experiments with different modality combinations are conducted to evaluate the individual effects of 1D SMILES semantics, 2D molecular topology, and 3D geometric information, as well as their complementary contributions after fusion. Second, learned gating weights reveal task-dependent modality preferences. Finally, t-SNE visualization is used to compare the distribution of molecular embeddings under different modality combinations, thereby evaluating the discriminative capability of fused representations in terms of intra-class compactness and inter-class separability. These analyses assess LSP modality complementarity and the structural consistency and task discriminability of fused representations. The corresponding results are shown in Fig. 3 and Fig. 4.

**Fig. 3.**
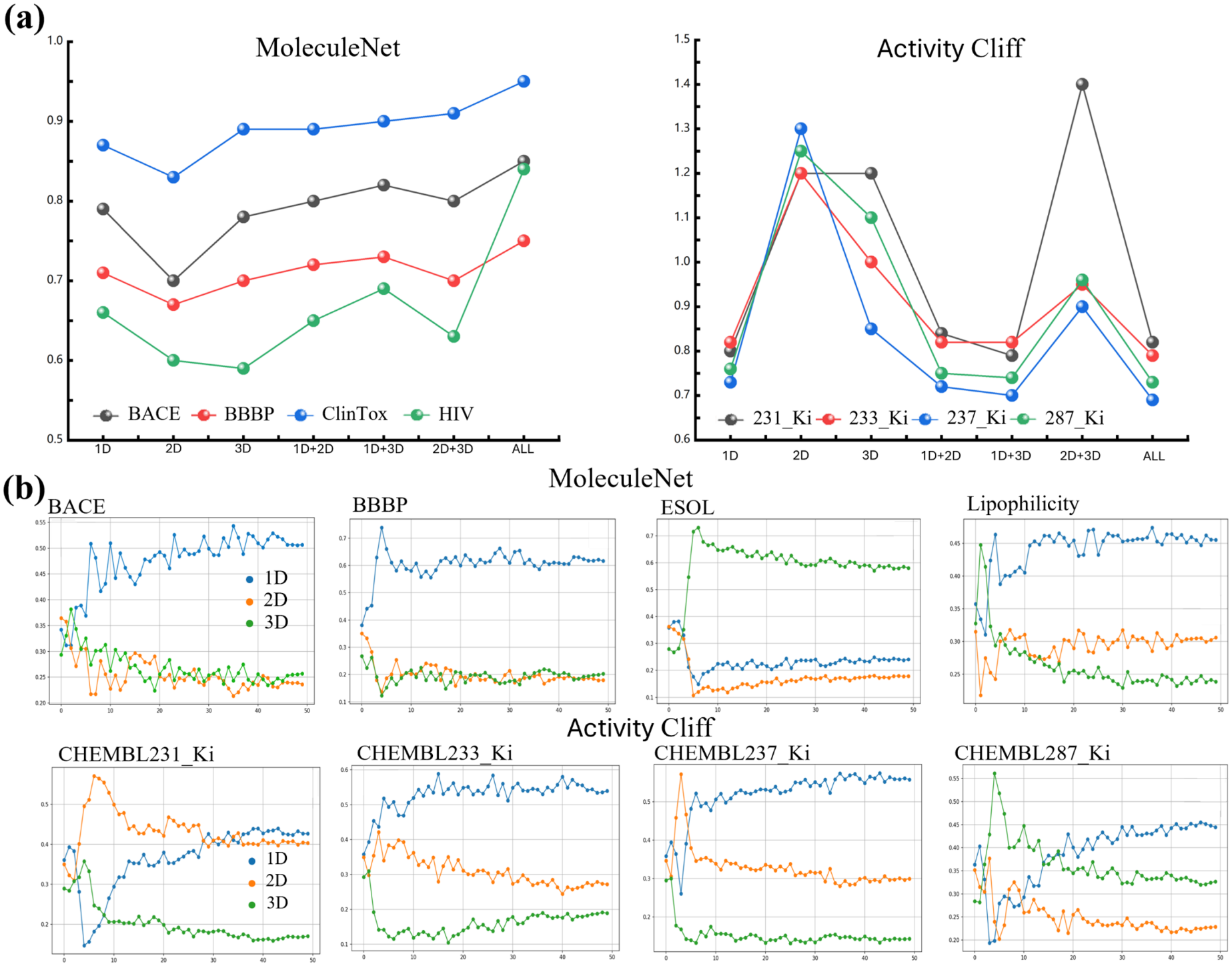
LSP information complementarity and dynamic fusion analysis. (a) Ablation results of different modality combinations on MoleculeNet classification and Activity Cliff tasks. (b) Dynamic modality weights of 1D, 2D, and 3D representations during training. Here, 1D, 2D, and 3D denote SMILES semantics, molecular topology, and three-dimensional geometric information, respectively, while ALL denotes the joint use of all three modalities. MoleculeNet classification performance is measured by ROC-AUC (higher is better), whereas Activity Cliff regression performance is measured by RMSE (lower is better).

**Fig. 4.**
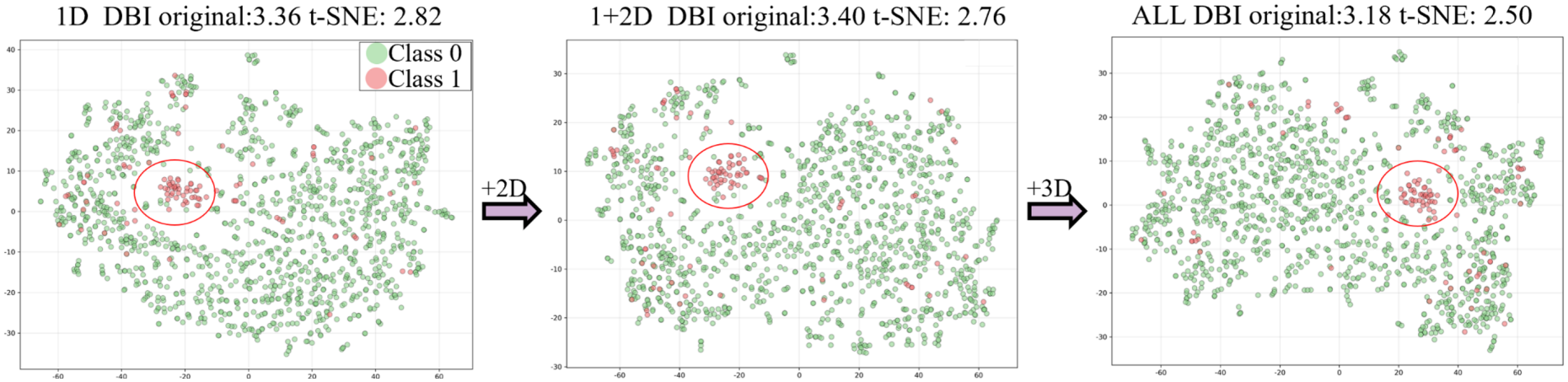
Representation-space analysis on the ClinTox dataset. t-SNE visualizations are shown for the 1D, 1D+2D, and ALL modality settings. DBI denotes the Davies-Bouldin index, for which a lower value indicates better clustering compactness and separation.

Fig. 3a summarizes the ablation results of different modality combinations on MoleculeNet classification and Activity Cliff tasks. The results show that different single modalities exhibit task-dependent performance, indicating that 1D SMILES semantics, 2D molecular topology, and 3D geometric information capture different aspects of molecular structures. On MoleculeNet tasks, the 3D modality performs better on ClinTox, suggesting its importance for toxicity-related properties, while the 1D modality shows relatively strong performance on BACE and BBBP, reflecting the contribution of chemical semantic information to bioactivity and blood-brain barrier permeability prediction. Although the 2D single modality performs relatively weaker overall, its combination with 1D or 3D representations improves performance, demonstrating the complementary role of molecular topology. On Activity Cliff tasks, 1D semantics and 3D geometry achieve lower RMSE values on different ChEMBL subtasks, indicating that both semantic and geometric information are important for predicting fine-grained activity differences. Overall, most dual-modality combinations and the ALL setting achieve more stable performance, validating the effectiveness of collaborative LSP information modeling.

Fig. 3b shows the dynamic modality weights assigned by the gating network during training. The weights vary across tasks, indicating that Distill-Mol does not rely on fixed fusion weights but adaptively adjusts the contributions of semantic, topological, and geometric information according to task requirements. On MoleculeNet tasks, BACE, BBBP, and HIV rely more on the 1D semantic modality in the later training stage, whereas ClinTox assigns relatively higher weights to the 3D modality, suggesting that geometric information provides useful complementary cues for toxicity-related prediction. For Activity Cliff tasks, different ChEMBL subtasks exhibit distinct weight distributions, further indicating that fine-grained activity changes require task-aware balancing of multi-source molecular information. These results demonstrate the flexibility and effectiveness of the proposed dynamic fusion mechanism.

Fig. 4 presents the t-SNE visualization of molecular embeddings on the ClinTox dataset under different modality combinations. The DBI measured in the t-SNE space decreases from 2.82 for the 1D representation to 2.76 after adding 2D topological information and further decreases to 2.50 under the ALL setting. This progression indicates that topology provides complementary structural information and that the further incorporation of 3D geometry improves the compactness and separation of the embedding space. Although some samples from different classes remain overlapped, the ALL representation shows the most coherent overall clustering pattern. These results demonstrate that joint semantic, topological, and geometric modeling improves the organization and discriminative capability of molecular representations.

### 4.4. Cross-Modal Fusion and Complementarity Analysis

To understand the predictive behavior of the model and its decision-making basis, we conduct atom-level attention interpretability analysis on the BBBP dataset. This experiment not only characterizes how different modalities focus on molecular structures but also reveals how such attention affects the final prediction outcomes. Because the BBBP task is sensitive to molecular polarity distribution, topological structure, and three-dimensional conformation, we select three representative molecules. The modality-specific attention distributions of these molecules are then visualized to examine the complementary roles of the different modalities in molecular representation learning. The corresponding results are shown in Fig. 5.

**Fig. 5.**
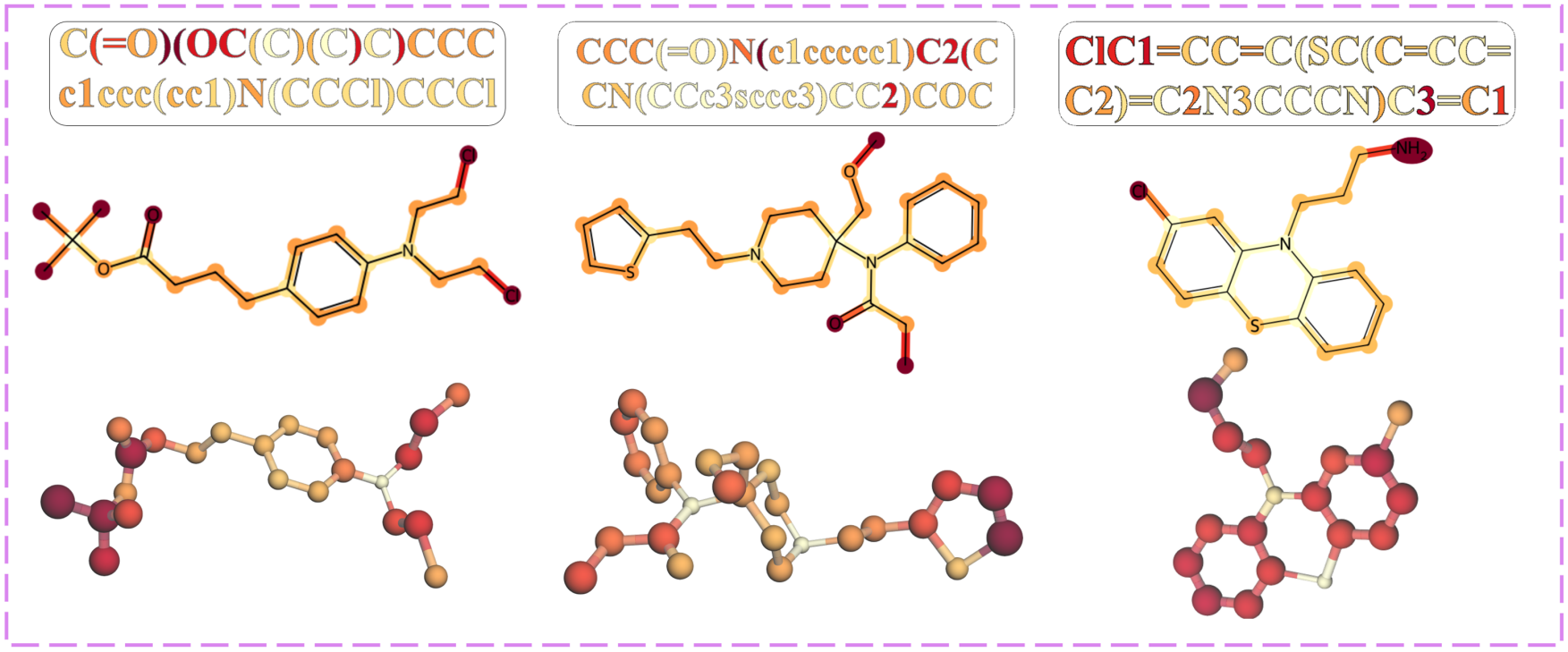
Multimodal attention complementarity analysis. Attention heatmaps are visualized for representative BBBP molecules under the 1D, 2D, and 3D modalities, corresponding to SMILES token-level attention, graph-based atom/substructure attention, and conformation-based spatial attention, respectively.

As shown in Fig. 5 the three modalities exhibit distinct yet complementary attention patterns on representative BBBP molecules, suggesting that Distill-Mol extracts discriminative cues from different representation spaces rather than repeatedly attending to the same structural regions. The 1D modality mainly focuses on chemically informative SMILES tokens and functional fragments, such as heteroatoms, halogen substitutions, carbonyl groups, aromatic moieties, and nitrogen-containing heterocycles, which are closely related to polarity, hydrophobicity, aromaticity, and hydrogen-bonding capacity. The 2D modality emphasizes graph-based topological substructures, including aromatic rings, heterocycles, fused-ring scaffolds, branch connections, and local connectivity around polar atoms, indicating its ability to capture atom-atom connectivity, substituent positions, and local structural environments. In contrast, the 3D modality highlights spatially informative regions determined by molecular conformation, such as exposed polar groups, terminal substituents, flexible chain segments, and spatially proximate atoms that may not be directly connected in the molecular graph. These 3D cues provide conformational and spatial information related to molecular shape, exposure of functional groups, and local geometric arrangement.

Notably, the attention maps show partial overlap on key chemical regions while preserving modality-specific focuses, reflecting the coordinated roles of chemical semantics, molecular topology, and spatial geometry. This indicates that the three modalities are not simply redundant, but jointly provide complementary information for BBBP prediction. Overall, these results show that Distill-Mol learns chemically meaningful and structurally complementary attention patterns, thereby enhancing the interpretability and discriminative capability of molecular property prediction.

### 4.5. SHAP-Based Interpretability Analysis

To further investigate the global decision-making patterns of Distill-Mol across different molecular property prediction tasks, we perform SHAP interpretability analysis on the BBBP, BACE, and ClinTox datasets [47]. Since Distill-Mol makes predictions based on high-dimensional multimodal representations that integrate SMILES semantics, 2D topology, and 3D geometry, directly interpreting latent features may not provide explicit chemical meanings. To improve interpretability and chemical relevance, we adopt a post-hoc surrogate modeling strategy. Specifically, the trained Distill-Mol model is first used to generate prediction outputs, while each molecule is mapped to a set of physicochemical descriptors with clear chemical semantics, including polarity, hydrophobicity, molecular size, ring count, and hydrogen-bond donor/acceptor features. These descriptors are then used as input features, and the predictions of Distill-Mol are used as supervision signals to train a random forest surrogate model that approximates the decision behavior of Distill-Mol in an interpretable feature space. Based on this surrogate model, SHAP values are computed to quantify the relative contributions of different physicochemical properties to model predictions. The SHAP value distributions, mean signed feature contributions, and feature correlation heatmaps for different tasks are shown in Fig. 6.

**Fig. 6.**
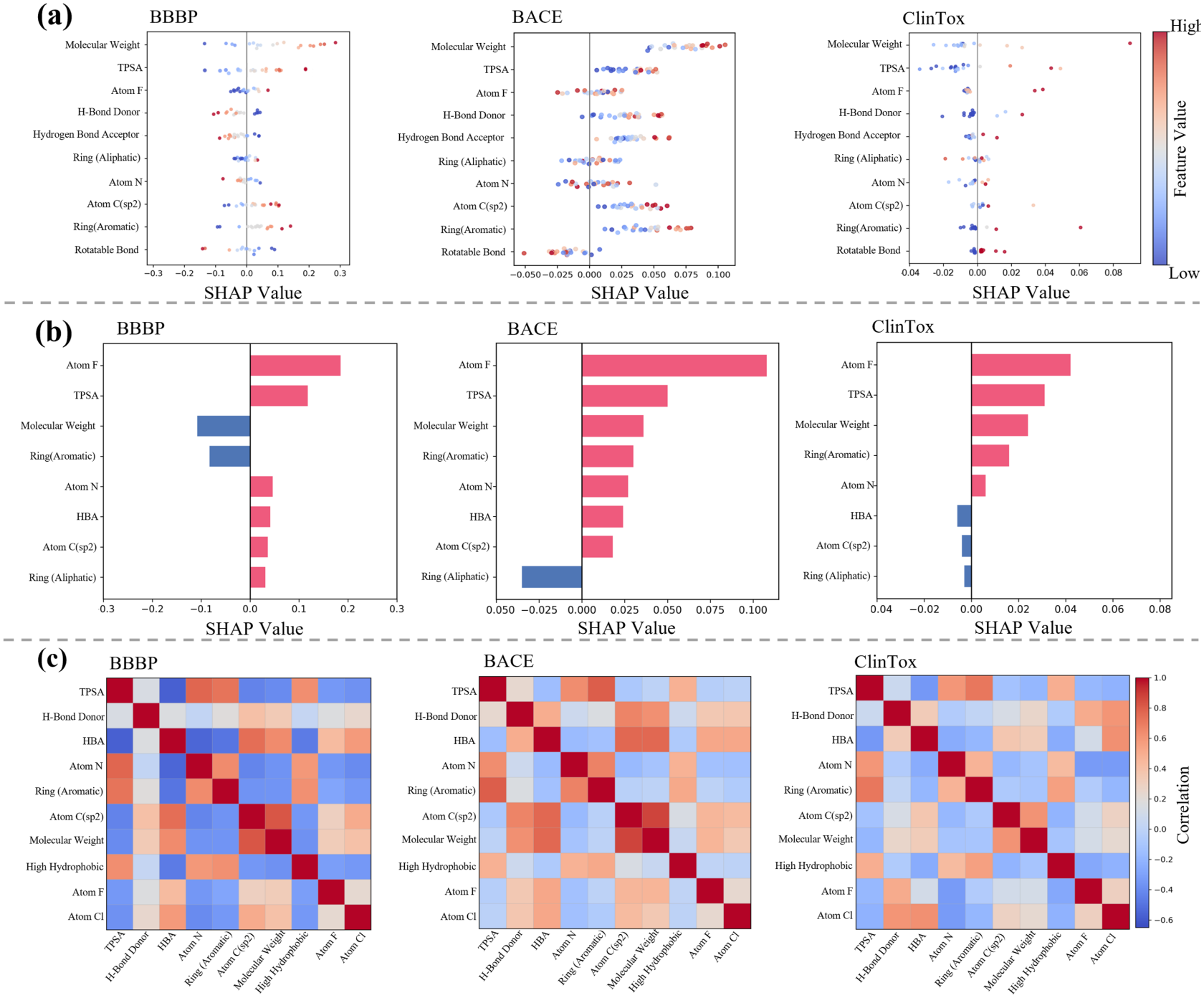
SHAP-based physicochemical interpretability analysis. (a) SHAP value distributions of molecular descriptors across samples. (b) Mean signed SHAP values of molecular descriptors across samples. (c) Correlation heatmaps among physicochemical descriptors.

As shown in Fig. 6a the SHAP value distributions reveal that feature contributions vary across samples and tasks, indicating that Distill-Mol does not rely on a single descriptor but integrates multiple physicochemical cues for prediction. For BBBP, hydrophobicity, TPSA, and hydrogen-bond-related features show substantial contributions, which is consistent with the known influence of lipophilicity and polarity on blood-brain barrier permeability. For BACE, aromatic rings, hydrophobicity, and hydrogen-bonding features are more prominent, suggesting that β-secretase inhibition prediction is closely related to hydrophobic scaffolds and local polar interactions. In ClinTox, high-contribution features are more dispersed, indicating that toxicity prediction is influenced by multiple factors, including polar functional groups, halogen substitutions, and molecular complexity. The mean signed SHAP values in Fig. 6b summarize the average direction of descriptor contributions across samples. Fluorine-related features and TPSA show positive mean contributions across the three tasks, whereas molecular weight and aromatic ring count show negative mean contributions on BBBP and positive mean contributions on BACE and ClinTox. These averages describe net contributions rather than global feature importance, since positive and negative attributions can cancel out. In addition, the correlation heatmaps in Fig. 6c reveal meaningful relationships among descriptors, including clusters of polarity-related features and hydrophobic/aromatic features. These results suggest that Distill-Mol’s predictions are chemically consistent in the interpretable physicochemical feature space. Overall, the SHAP analysis indicates that the multimodal representations learned by Distill-Mol not only improve predictive performance but also capture task-relevant structure-property relationships.

### 4.6. Validation of CAN/CGN Perturbations

To verify whether the proposed conformation angle noise (CAN) and coordinate Gaussian noise (CGN) generate reasonable and complementary three-dimensional perturbations, we analyze the energy distributions of perturbed conformations on both classification and regression datasets. Specifically, we compare the energy variations induced by different perturbation strategies with the low-energy threshold (LET), aiming to examine whether CGN preserves locally stable low-energy perturbations and whether CAN covers broader conformation-level variations. The perturbation energy distributions on classification and regression tasks are shown in Fig. 7 and Fig. 8 respectively.

**Fig. 7.**
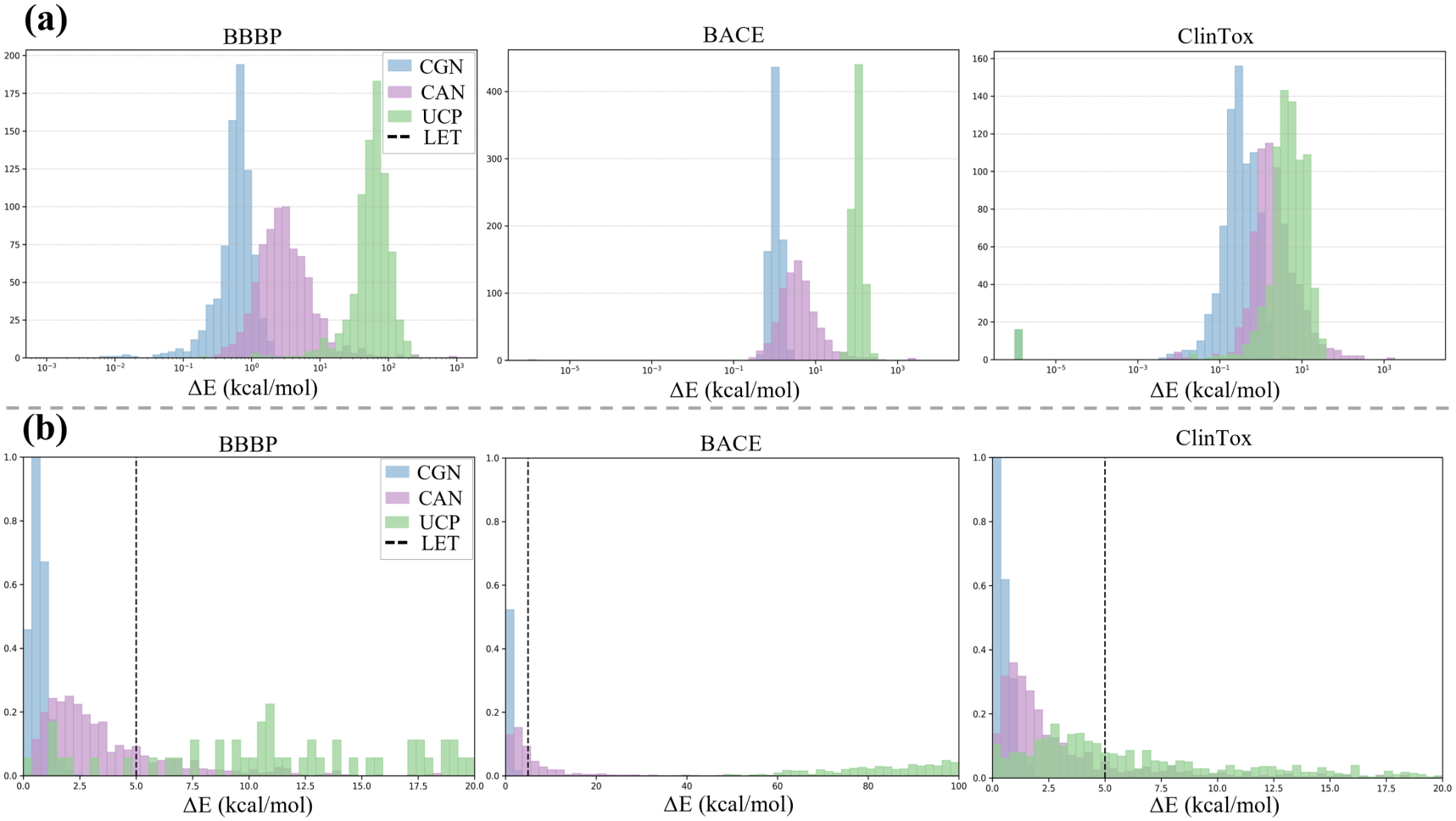
Perturbation energy distribution analysis on classification datasets. (a) Overall energy distributions of conformations generated by CGN, CAN, and UCP. (b) Detailed distributions within the low-energy region. CGN denotes coordinate Gaussian noise, CAN denotes conformation angle noise, UCP denotes unconstrained coordinate perturbation, and the dashed LET line denotes the low-energy threshold.

**Fig. 8.**
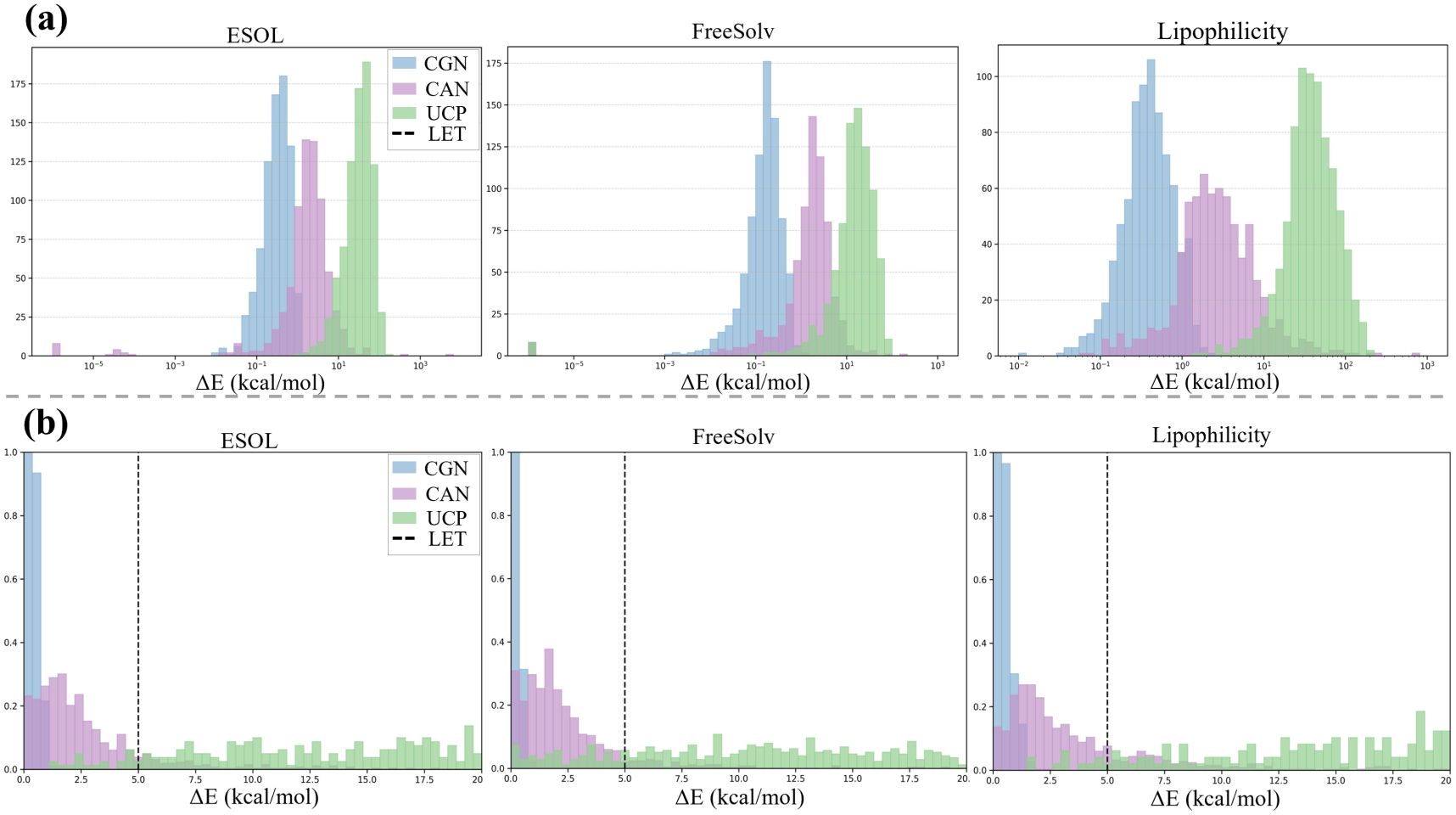
Perturbation energy distribution analysis on regression datasets. (a) Overall energy distributions of different perturbation strategies. (b) Detailed distributions within the low-energy region. The meanings of CGN, CAN, UCP, and LET are the same as in Fig. 7.

As shown in Fig. 7 and Fig. 8 conformations generated by different perturbation strategies exhibit distinct energy distribution patterns. CGN-generated conformations are mainly concentrated in the low-energy region, indicating that CGN introduces relatively small local perturbations around the original conformation. This property makes CGN suitable for the student model, which is expected to learn robust geometric representations under locally stable perturbations. In contrast, CAN produces a broader energy distribution by perturbing torsion angles around rotatable single bonds, thereby covering larger conformation-level variations. This makes CAN suitable for the teacher model, which provides richer conformational geometric knowledge for distillation. Compared with CGN and CAN, UCP shows more dispersed energy distributions, with some samples deviating substantially from the low-energy region, suggesting that unconstrained coordinate perturbations may introduce unstable or chemically unreasonable conformations.

The above trends are consistent across both classification and regression datasets, indicating that the proposed CAN/CGN perturbation design is stable under different task settings. The detailed views of the low-energy region further show that CGN perturbations are concentrated near the LET, whereas CAN extends to relatively higher-energy regions and provides greater conformational diversity. Therefore, CGN and CAN are complementary in terms of perturbation scale and energy distribution: CGN emphasizes local low-energy stability, while CAN captures broader conformation-level variations. This combination provides reasonable geometric supervision for teacher-student distillation and helps Distill-Mol learn 3D molecular representations with both local robustness and conformational sensitivity.

## 5. Conclusion

Molecular representation learning is a fundamental problem in computational chemistry, drug discovery, and materials design, aiming to rapidly and accurately evaluate the physicochemical properties, biological activities, and safety profiles of candidate molecules while reducing reliance on costly experimental measurements and quantum-chemical calculations. From the perspective of language-structure-physics (LSP) collaborative modeling, this paper proposes Distill-Mol, a multimodal molecular representation learning framework based on cross-modal fusion and hierarchical geometric distillation. Specifically, Distill-Mol uses SMILES sequences, 2D molecular graphs, and 3D conformations to model molecular chemical semantics, topological structures, and spatial geometric information, respectively, and maps these heterogeneous modalities into a unified representation space to enhance the consistency and complementarity of multi-level molecular information. Furthermore, a geometry-aware teacher-student distillation mechanism is designed under CAN/CGN perturbations, where CAN captures conformation-level variations induced by rotatable single bonds and CGN models local coordinate perturbations, thereby enabling cross-scale geometric knowledge transfer. In addition, the task-aware cross-modal fusion strategy adaptively adjusts the contribution of each modality according to different property prediction tasks, improving the flexibility and task adaptability of molecular representation learning.

Experimental results on benchmark datasets such as MoleculeNet and QM9 show that Distill-Mol achieves superior or competitive performance across multiple classification and regression tasks, while supplementary DTA and DTI experiments further evaluate the transferability of its molecular representations to biomedical interaction modeling scenarios. Ablation studies, modality weight analysis, geometric perturbation validation, and interpretability analysis further demonstrate that LSP multimodal collaborative modeling, CAN/CGN-based geometric distillation, and dynamic cross-modal fusion all contribute to improved predictive performance and representation quality. These results indicate that integrating molecular semantics, topology, and geometric priors is beneficial for learning robust and task-adaptive molecular representations. Nevertheless, the current framework still has limitations in modeling realistic conformational ensembles, fine-grained electronic structures, and complex environmental effects. In future work, we will further incorporate physicochemical priors such as multi-conformation sampling, local charges, electronic structures, molecular force fields, and solvent effects. We will also extend Distill-Mol to more complex biomedical scenarios, including drug-drug interaction prediction, protein-ligand binding site modeling, and ADMET-related risk assessment, thereby enhancing its applicability to real-world drug discovery and materials design.

## Appendix A. Dataset Details

**Table A.1.** MoleculeNet dataset statistics and task settings.

| Dataset | #Molecules | #Tasks | Task type | Metric |
|---|---|---|---|---|
| BACE | 1513 | 1 | Inhibitor prediction | ROC-AUC |
| BBBP | 2039 | 1 | Blood-brain barrier permeability | ROC-AUC |
| ClinTox | 1478 | 2 | Clinical toxicity prediction | ROC-AUC |
| Tox21 | 7831 | 12 | Toxicity prediction | ROC-AUC |
| ToxCast | 8477 | 617 | Toxicity assay prediction | ROC-AUC |
| MUV | 93087 | 17 | Virtual screening prediction | ROC-AUC |
| SIDER | 1427 | 27 | Side-effect prediction | ROC-AUC |
| HIV | 41127 | 1 | Anti-HIV activity prediction | ROC-AUC |
| ESOL | 1128 | 1 | Aqueous solubility prediction | RMSE |
| FreeSolv | 642 | 1 | Hydration free energy prediction | RMSE |
| Lipophilicity | 4200 | 1 | Lipophilicity prediction | RMSE |

### A.1. MoleculeNet Dataset Details

The MoleculeNet benchmark covers a wide range of molecular property prediction tasks, including classification tasks and regression tasks. In this work, the classification datasets include BACE, BBBP, ClinTox, Tox21, ToxCast, MUV, SIDER, and HIV, while the regression datasets include ESOL, FreeSolv, and Lipophilicity. These datasets correspond to different molecular properties, including bioactivity, blood-brain barrier permeability, toxicity, side effects, antiviral activity, solubility, hydration free energy, and lipophilicity. Therefore, they provide a comprehensive benchmark for evaluating the predictive performance and generalization ability of molecular representation learning models across different task types and data distributions. We follow standard data splitting and evaluation protocols, and the reported results are averaged over ten independent runs for all tasks.

### A.2. QM9 Dataset Details

The QM9 dataset contains approximately 130,000 small molecules and their corresponding quantum-chemical properties. It is widely used to evaluate the ability of molecular models to capture three-dimensional geometry and electronic properties. Each molecule contains up to nine heavy atoms selected from C, O, N, and F, and multiple physicochemical attributes are computed using density functional theory (DFT). Therefore, QM9 provides a suitable benchmark for evaluating geometric modeling ability and physical consistency in molecular representation learning.

In this work, we select commonly used quantum-chemical prediction tasks, including dipole moment (*µ*), polarizability (*α*), highest occupied molecular orbital energy (HOMO), lowest unoccupied molecular orbital energy (LUMO), energy gap (Gap), zero-point vibrational energy (ZPVE), and thermodynamic energies (*U*_0_, *U*, and *H*). These properties are highly sensitive to molecular three-dimensional geometry and electronic distribution, and thus can effectively evaluate the geometric modeling capability of the proposed method. All QM9 experiments are conducted under a unified data splitting and evaluation protocol, and mean absolute error (MAE) is adopted as the evaluation metric.

## APPENDIX B. Evaluation Metrics

Different evaluation metrics are adopted for different task types. ROC-AUC is used for classification tasks, while RMSE and MAE are used for regression tasks.

ROC-AUC measures the ranking ability of a classification model for positive and negative samples and is defined as

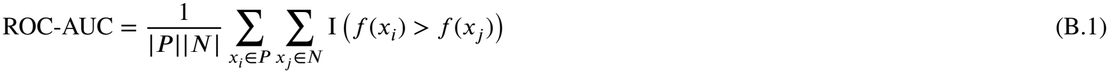

where *P* and *N* denote the sets of positive and negative samples, respectively, *f*(*x*) denotes the predicted score of sample *x*, and I(*⋅*) is the indicator function. This metric can be interpreted as the probability that a randomly selected positive sample receives a higher score than a randomly selected negative sample.

RMSE measures the root mean squared prediction error and is defined as

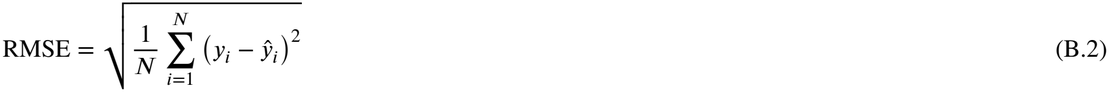

where *y*_*i*_ and ŷ_*i*_ denote the ground-truth value and predicted value of the *i*-th sample, respectively, and *N* is the number of samples. RMSE is more sensitive to large prediction errors.

MAE measures the average absolute prediction error and is defined as

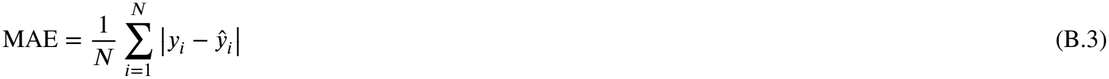

Compared with RMSE, MAE is less sensitive to outliers and more directly reflects the overall prediction deviation.

## Data availability

All datasets used in this study are publicly available from their original sources, which are cited in the manuscript. The source code for Distill-Mol is available at https://github.com/ano-molresearch/distill-mol.

